# Towards an optimised processing pipeline for diffusion MRI data: Effects of artefact corrections on diffusion metrics and their age associations in UK Biobank

**DOI:** 10.1101/511964

**Authors:** Ivan I. Maximov, Dag Alnæs, Lars T. Westlye

## Abstract

Increasing interest in the structural and functional organization of the human brain in health and disease encourages the acquisition of big datasets consisting of multiple neuroimaging modalities accompanied by additional information obtained from health records, cognitive tests, biomarkers and genotypes. Diffusion weighted magnetic resonance imaging data enables a range of promising imaging phenotypes probing structural connections as well as macroanatomical and microstructural properties of the brain. The reliability and biological sensitivity and specificity of diffusion data depend on processing pipeline. A state-of-the-art framework for data processing facilitates cross-study harmonisation and reduces pipeline-related variability. Using diffusion data from the UK Biobank we provide a comprehensive evaluation of different processing steps that have been suggested to reduce imaging artefacts and improve reliability of diffusion metrics. We consider a general pipeline comprising 7 post-processing blocks: noise correction; Gibbs ringing correction; evaluation of field distortions; susceptibility, eddy-current and motion-induced distortion corrections; bias field correction; spatial smoothing; and final diffusion metric estimations. Based on this evaluation, we suggest an optimised diffusion pipeline for processing of diffusion weighted imaging data.

## Introduction

Increasing interest in the role of individual differences in human brain architecture in health and disease has stimulated the neuroscience community to initiate a number of large brain data projects. Due to the attractive combination of increasing availability, low costs, its non-invasive nature and high sensitivity magnetic resonance imaging (MRI) including T1/T2-weighted images, functional MRI with tasks and resting state, perfusion and diffusion weighted imaging has become the preferred and standard brain imaging modality in these large efforts, including the UK Biobank (UKB) (Miller et al., 2016). Brain imaging data are often accompanied by clinical and biological information such as psychological tests, blood samples, genetic data and other parameters depending on original project targets. Combining all obtained information into a common statistical framework benefits from universally adapted post-processing pipelines for harmonised data quality assessment and manipulation.

Diffusion MRI is based on the effect of the Brownian motion of water molecules in biological tissue (Basser et al., 1994) and allows one to probe and visualise brain organisation at the micrometer scale (Johansen-Berg and Behrens, 2014). Current impetuous growth of theoretical and experimental diffusion MRI approaches (Novikov et al., 2018) has offered various diffusion models and sequences in order to effectively describe the signal decay due to water diffusion. Advanced diffusion measurements are technically challenging and optimal data quality places high demands on practical implementation and protocol, including hardware gradient system and coil. Due to limited time and technical constraints researchers designing imaging studies face various trade-offs, influencing, e.g. signal-to-noise ratio (SNR) and options related to the specific pulse sequences such as mono-or bipolar diffusion gradients etc.

Before diffusion metric estimations and statistical analysis there are many different approaches of quality control (QC) and corrections applied to the data in order to verify diffusion data integrity (Alfaro-Almagro et al., 2018; Farzinfar et al., 2013; Esteban et al., 2017; Hasan, 2007; Oguz et al., 2014). Ideally, the QC methods should rapidly identify and correct typical artefacts such as head motion, discarded volumes, and low SNR, which may be particularly present at high diffusion weightings, also known as *b*-values. Despite recent major developments and improvements (Alfaro-Almagro et al., 2018; Ciu et al., 2013; Miller et al., 2016; Roalf et al., 2016), automated procedures for QC and artefact reduction largely represent unresolved challenges in the imaging community.

Various post-processing steps have been suggested in order to correct common sources of noise and distortions, including thermal noise evaluation (Veraart et al., 2016a; Veraart et al., 2016b), Gibbs ringing correction (Kellner et al., 2016; Veraart et al., 2016c), susceptibility distortion correction (Andersson and Sotiropoulos, 2016a), motion correction (Andersson and Sotiropoulos, 2016a; Andersson et al., 2016b), correction of physiological noise and outliers (Maximov et al., 2011; Maximov et al., 2015; Sairanen et al., 2018; Walker et al., 2011), and eddy current induced geometrical distortions (Taylor et al., 2016). However, although the application of even part of the post-processing steps such as noise correction has been demonstrated to improve sensitivity and provide additional information about absolute diffusion metrics (Kochunov et al., 2018), systematic evaluations of the effects of the different steps on the diffusion metrics are scarce.

Several minimal post-processing pipelines have been recommended in order to prepare structural, functional and diffusion MRI data (Alfaro-Almagro et al., 2018; Ciu et al., 2013; Glasser et al., 2013; Sotiropoulos et al., 2013). Albeit fairly comprehensive, none of the recommended protocols include all steps listed above. For example, the UKB diffusion pipeline first employs fieldmap generation using the anterior-posterior (AP) and posterior-anterior (PA) images of original diffusion data. In turn, the selection of most reliable AP-PA images is performed by an estimation of relative correlations over all AP-PA images in order to find the most accurate reference image.

Thus, the UKB data exhibit only one diffusion-specific pipeline step based on *eddy* (Andersson et al., 2016a; Andersson et al., 2016b; Andersson et al., 2017), correcting the eddy currents and head motion, susceptibility artefacts and identification and replacement of outlier slices.

With the aim to identify the most efficient and adequate pipeline for diffusion data analysis we tested the effects of various diffusion post-processing steps on different diffusion scalar metrics, based on diffusion tensor imaging (Basser et al., 1994), diffusion kurtosis imaging (Jensen et al., 2005), and white matter tract integrity (Fieremans et al., 2011) using UKB data. In order to assess to which degree the chosen pipeline influences across-subject analysis and corresponding biological interpretations we compared estimated age-curves (Grinberg et al., 2017; Tamnes et al., 2017; Westlye et al., 2010; Westlye et al., 2012) of the diffusion metrics between pipelines using tract-based spatial statistic (Smith et al., 2006, 2007).

## Methods and Materials

### Subjects and data

Table 1 summarises the demographics of the 218 UKB subjects included in the present work. We computed diffusion scalar metrics using four different pipelines, i.e. the total number of datasets included in the analysis is 872. An accurate overview of the UKB data acquisition, protocol parameters, and image validation can be found elsewhere (Alfaro-Almagro et al., 2018; Miller et al., 2016). The original UKB post-processing pipeline is described in details online (http://biobank.ctsu.ox.ac.uk/crystal/docs/brain_mri.pdf).

**Table 1.** Demographic data of the used UK Biobank sample.

| Subgroups (years) | Number of subjects | Age (Mean/std) years | Sex (F/M) |
| --- | --- | --- | --- |
| “40” | 12 | 40.40/0.08 | 6/6 |
| “42” | 13 | 42.08/0.29 | 6/7 |
| “44” | 16 | 43.92/0.28 | 8/8 |
| “46” | 14 | 46.01/0.31 | 7/7 |
| “48” | 13 | 48.00/0.32 | 7/6 |
| “50” | 15 | 50.06/0.29 | 8/7 |
| “52” | 15 | 52.07/0.27 | 7/8 |
| “54” | 11 | 54.11/0.35 | 7/4 |
| “56” | 15 | 55.97/0.26 | 7/8 |
| “58” | 13 | 57.98/0.29 | 6/7 |
| “60” | 15 | 59.99/0.27 | 8/7 |
| “62” | 14 | 61.95/0.26 | 7/7 |
| “64” | 12 | 64.11/0.25 | 7/5 |
| “66” | 14 | 66.11/0.25 | 8/6 |
| “68” | 14 | 68.08/0.27 | 7/7 |
| “70” | 12 | 69.81/0.18 | 6/6 |
| Total | 218 | 54.95/9.09 | 112/106 |

The pipeline used in the present work is shown in Figure 1. We divided the post-processing flow into 7 general blocks. Additional block *i* (marked by blue frame in Fig.1) consists of extra steps allowing one to substitute or extend used algorithms. An advantage of the discussed pipeline is freely accessible open source code for all processing steps. Below we shortly describe each step in the suggested order.

**Figure 1.**
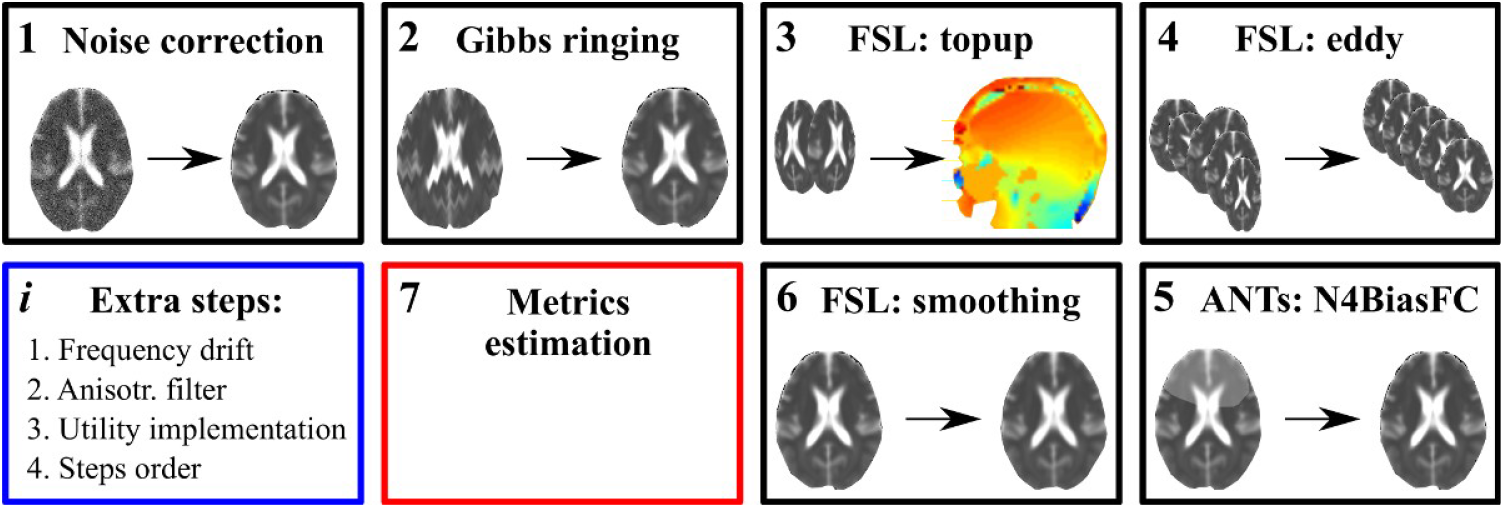
Schematic representation of a general pipeline. Numbers in the upper left corner correspond to the step order. The step 7 is an estimation of final diffusion metrics depending on used diffusion model. The step *i* presents a possible variability in the pipeline but omitted in the present work, for example, a frequency drift correction, application of different spatial filters (isotropic vs anisotropic), difference in algorithmic utility implementations (ANTs vs FSL), permutations in the step orders (step 5 vs step 4).

#### 1. Noise correction

The noise in diffusion data is spatially dependent in the case of multi-channel receive coils (Aja-Fernandez et al., 2014; Andre et al., 2014; Maximov et al., 2012). Principle component analysis of Marchenko-Pastur noise-only distribution provides an accurate and fast method of noise evaluation (Veraart et al., 2016a; Veraart et al., 2016b), thereby enabling signal-to-noise ratio enhancements by the Rician noise correction. In the presented work we used the original Veraart’s Matlab code (The Mathworks, Natick, Massachussets, USA): https://github.com/NYU-DiffusionMRI/mppca_denoise. The noise correction methods are regularly improved and in the future it might be substituted by a more efficient approach (see an evaluation of methods, for example, in reviews of Aja-Fernandez et al., 2014; Manjon et al., 2015).

#### 2. Gibbs-ringing correction

Various artefacts appearing in the raw data due to table vibration (Gallichan et al., 2010), radiofrequency based distortions, incorrect magnetic field gradient calibration (McRobbie et al., 2006) can significantly degrade the diffusion data. One of the most frequent artefact is known as the Gibbs ringing artifact. This appears due to a A>space truncation along finite image sampling and can be suppressed by *post hoc* methods (Kellner et al., 2016; Perrone et al., 2015; Veraart et al., 2016c). Here we used the approach developed by Kellner and colleagues (Kellner et al., 2016) and the original Matlab code: https://bitbucket.org/reisert/unring.

#### 3. EPI distortions

Diffusion data acquisition is based on echo-planar imaging (EPI), which is susceptible to multiple distortions originating from a magnetic field inhomogeneity. A few approaches have been developed to correct field inhomogeneities: a simple and robust method based on field mapping; a method based on evaluation of point spread function; and reversed gradient approach (Wu et al., 2008). *FSL* (Smith et al., 2004) offers an excellent utility for the EPI geometric distortion correction *(topup,* https://fsl.fmrib.ox.ac.uk/fsl/fslwiki/topup, Andersson et al., 2003). *Topup* requires data with opposite phase-encoding directions for the non-diffusion weighted images (so called *b_0_* images), for example, anterior-posterior and posterior-anterior pair or left-right and right-left pair.

#### 4. Motion, eddy current and susceptibility distortion correction

*Topup* and *eddy* works together for correcting distortions appeared due to eddy currents, head motion and susceptibility originated artefacts. The GPU accelerated version of *eddy (eddy_cuda)* allows one to significantly speed up the computations as well as providing additional options such as in slice alignments, improved outlier detection and multi-band dataset estimations (https://fsl.fmrib.ox.ac.uk/fsl/fslwiki/eddy/UsersGuide) (Andersson et al., 2016a; Andersson et al., 2016b; Andersson et al., 2017).

#### 5. Field non-uniformity

MR images possess a low frequency intensity shift appearing as intensity inhomogeneity over the image. Several studies have evaluated its influence on the intra- and inter-subject reproducibility of T1-weighted structural MRI data (Banerjee and Maji, 2015; Ganzetti et al., 2016). However, less has been published regarding effects of non-uniformity correction on diffusion data. In order to avoid bias based on the field non-uniformity we applied a bias field correction for *b0* image. Then, the estimated field map was applied to all diffusion images in order to decrease the field inhomogeneity. We used the *N4BiasFieldCorrection* utility from ANTs (Tustison et al., 2010). An applied order of the bias field correction step is discussed below.

#### 6. Spatial smoothing

After all previous steps, the diffusion data, in theory, are ready for diffusion scalar metric evaluation. In order to increase SNR, which may be particularly beneficial for the numerical stability of advanced diffusion models (Maximov et al., 2017; Vellmer et al., 2018), we applied spatial smoothing of the raw diffusion data. For simplicity, we used isotropic smoothing with a Gaussian kernel 1 mm^3^ implemented in the FSL function *fslmaths.*

#### 7. Metric estimation

UKB diffusion data acquisition was done using a multi-shell protocol with *b* = 1000 and 2000 s/mm^2^ in addition to *b=0.* The dataset allows one to obtain typical diffusion metrics such as conventional diffusion tensor imaging (cDTI) (Basser et al., 1994), namely, fractional anisotropy (FA), mean, axial, and radial diffusivity (MD, AD, and RD, respectively); diffusion kurtosis imaging (DKI) (Jensen et al., 2005) with FA, MD, AD, RD, mean, axial, and radial kurtosis (MK, AK, RK, respectively); white matter tract integrity (WMTI) (Fieremans et al., 2011) metrics with axonal water fraction (AWF), extra-axonal axial and radial diffusivities (AE and RE), and tortuosity (Tort). These metrics are based on a cumulant expansion of the diffusion propagation function, i.e. strictly speaking they do not represent a comprehensive diffusion biophysical model (Novikov et al., 2018). Nevertheless, these maps are very popular and easy to obtain in clinical studies. For simplicity, we consider here only DTI, DKI and WMTI metrics. For DKI, we used an approach proposed by Veraart et al., (2013) and the original Matlab code (https://github.com/NYU-DiffusionMRI/Diffusion-Kurtosis-Imaging). DTI was estimated using *DTIFIT* in FSL, by means of a linear weighted least squares option in command line for the shell *b* = 1000 s/mm^2^. We assume that the original UKB DTI metrics were estimated with the same option, although it was not mentioned in the description (Miller et al., 2016).

##### i. Additional options

Some of the steps can be substituted by other approaches or implementations. For example, nonuniformity field corrections used in functional MRI and brain tissue segmentation may increase the accuracy of the motion correction (Ganzetti et al., 2016). The applied isotropic spatial filtering even with a quite small Gaussian kernel introduces blurring of tissue borders and increase partial voluming. A classical anisotropic diffusion filter based on the Perona-Malik algorithm (Perona and Malik, 1987) may provide an alternative with less blurring (Vellmer et al., 2018; Van Hecke et al., 2010). Therefore, we suggest to carefully consider the influence of different degradation factors on the diffusion image quality and to choose a reliable and robust tool for the correction step tailored to the study (see, for example, considerations related to neonatal neuroimaging: Bastiani et al., 2019).

### Temporal SNR

In order to quantify the difference between the pipelines on a conventional data quality metric we estimated temporal signal-to-noise ratios (tSNR, Roalf et al., 2016) for each pipeline, which allows one to present a single numeric metric characterising the whole brain diffusion weighted dataset and to perform comparative estimations of data quality (Tønnesen et al., 2018). In order to accommodate the estimation to multi-shell data we separated the datasets into two independent *b*-shells with *b* = 1000 and 2000 s/mm^2^, respectively. In each case, tSNR was evaluated for S4, S5, S7 and UKB pipelines (see below).

### Statistical analysis

In order to perform a statistical comparison between pipelines we used TBSS (Smith et al., 2006). Initially, all volumes were aligned to the FMRI58_FA template, supplied by FSL, using non-linear transformation realised by FNIRT (Andersson et al., 2007). Next, a mean FA image of all subjects was obtained and thinned in order to create mean FA skeleton. Afterwards, all FA images were thresholded at FA > 0.2. The maximal FA values were projected onto the skeleton in order to minimise confounding effects due to partial voluming.

Voxel-wise associations between age and the diffusion metrics were tested using general linear models (GLMs), including sex as covariate. The statistical analysis was performed using permutation-based inference implemented in *randomise* (Winkler et al., 2014) with 5000 permutations. Threshold-free cluster enhancement (TFCE, Smith and Nichols, 2009) was used. Statistical *p* value maps were thresholded at *p* < 0.05 corrected for multiple comparisons across space. In addition to voxel-wise statistics, we used diffusion metrics averaged over the skeletons for visualisation and plotting of differences between pipelines. The pipeline difference was visualised by correlation plots with estimated Pearson correlation coefficients *(corrplot* function of Matlab, The Mathworks, Natick, MA USA).

In order to assess to which degree the chosen pipeline can influence the interpretation of the results we compared the estimated age-related slopes for the diffusion metrics between S7 and UKB pipelines. Averaged diffusion metrics were used for plotting the age-curves. Linear regressions were performed using ordinary least squares fitting extracting regression slopes and intercepts. The Pearson correlation coefficient *r* and the coefficient of determination *r*^2^ between age and diffusion metrics were estimated using the *corr* function of Matlab.

## Results

### Voxel-wise comparisons

Figure 2 shows the scatter plots for the DTI metrics (FA, MD, AD, and RD) obtained from the DKI signal fitting and four pipelines. Left column of correlation plots corresponds to the mean values of diffusion metrics evaluated over the skeleton, the right column represents the standard deviations. Briefly, the results revealed high correlations of the diffusion metrics between all pipelines, however, the S7 pipeline exhibited significantly (Levene’s test with p < 0.001) reduced standard deviation (histogram peak values are at MD – 0.18; AD – 0.29; RD – 0.15) compared to the original UKB pipeline (histogram peak values are at MD – 0.21; AD – 0.35; RD – 0.23). Figure 3 shows the correlation plots for the DKI scalar metrics (MK, AK, and RK). The kurtosis scalar metrics were highly correlated between pipelines. The standard deviations are also lower in the S4, S5, and S7 pipelines compared to UKB. Figure 4 shows the correlation plots for the WMTI scalar metrics (AWF, AE, RE, and Tort). Since the estimation of WMTI metrics were based on the DKI values, the WMTI diffusion metrics exhibited quite high correlations for all pipelines similar to Fig. 3. The standard deviation of S7 pipeline was lower compared to all other pipelines (histogram peak values are at AWF – 0.07; AE – 0.40; RE – 0.18; Tort – 0.65). Figure 5 shows the scatter plots for the conventional single-shell (b = 1000 s/mm^2^) DTI metrics, suggesting similar relationships between the pipelines as for DKI, with lower standard deviation in the S7 pipeline compared to the original UKB pipeline.

**Figure 2.**
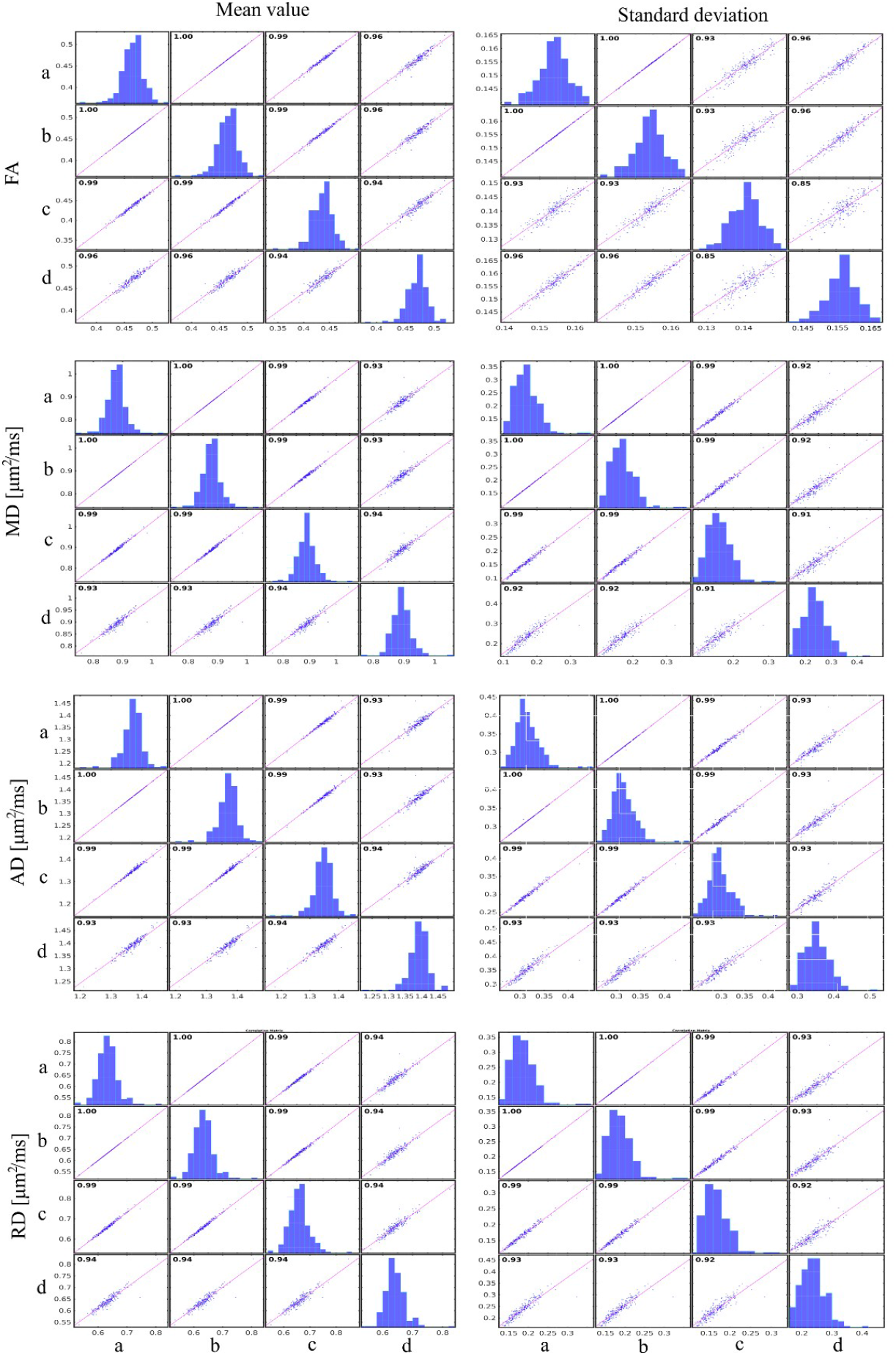
Correlation plots for diffusion metrics based on DKI fitting obtained for four different pipelines (see Fig. 1): a) up to Step 5; b) up to Step 4; c) up to Step 7; d) original UK Biobank pipeline. Diffusion metrics were averaged over estimated subject skeletons in the case of each pipeline in accordance with the TBSS preparation pipeline.

**Figure 3.**
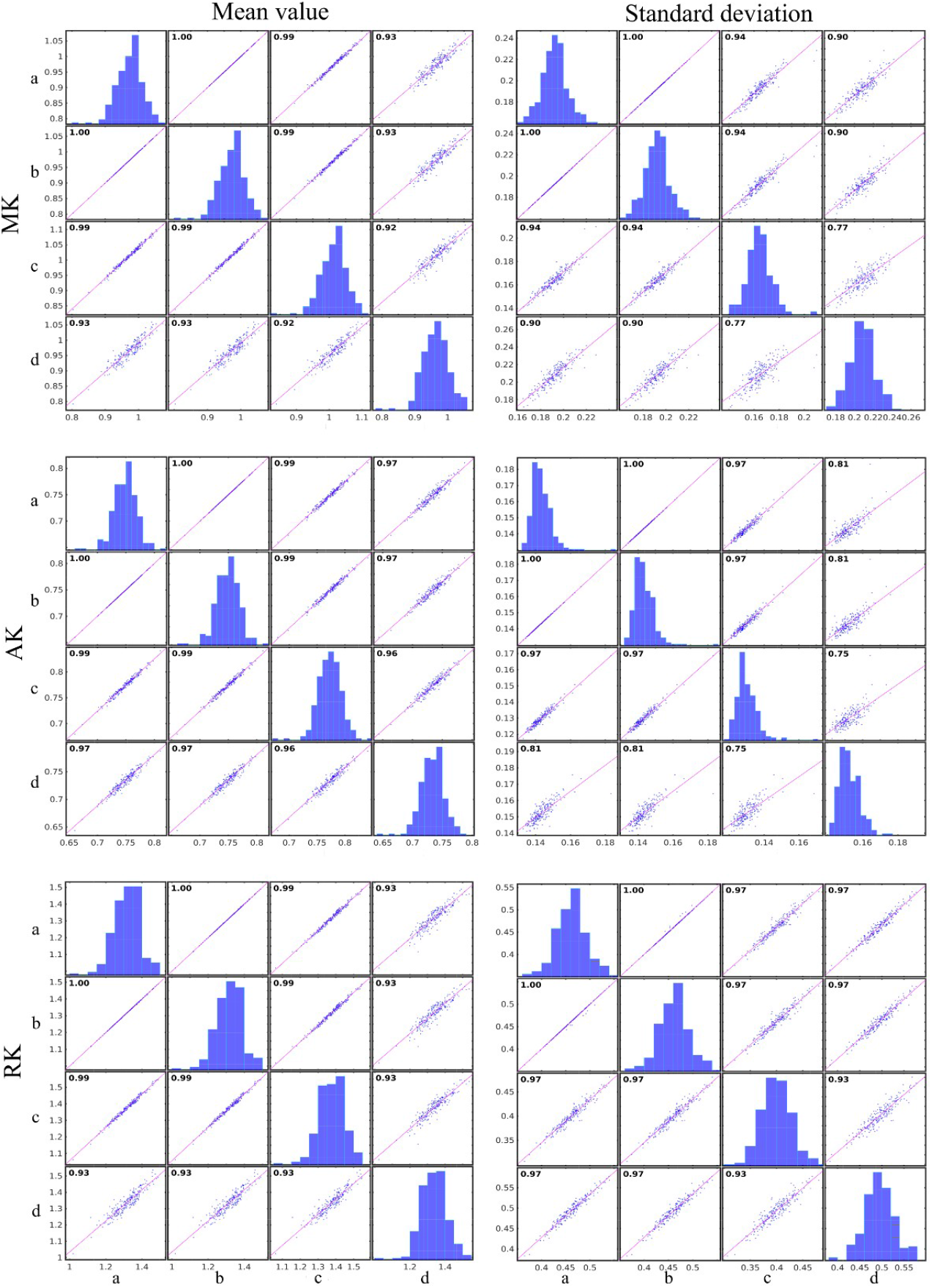
Correlation plots for diffusion metrics based on DKI fitting obtained for four different pipelines: a) up to Step 5; b) up to Step 4; c) up to Step 7; d) original UKB pipeline. Diffusion metrics were averaged over estimated subject skeletons in the case of each pipeline in accordance with the TBSS preparation pipeline.

**Figure 4.**
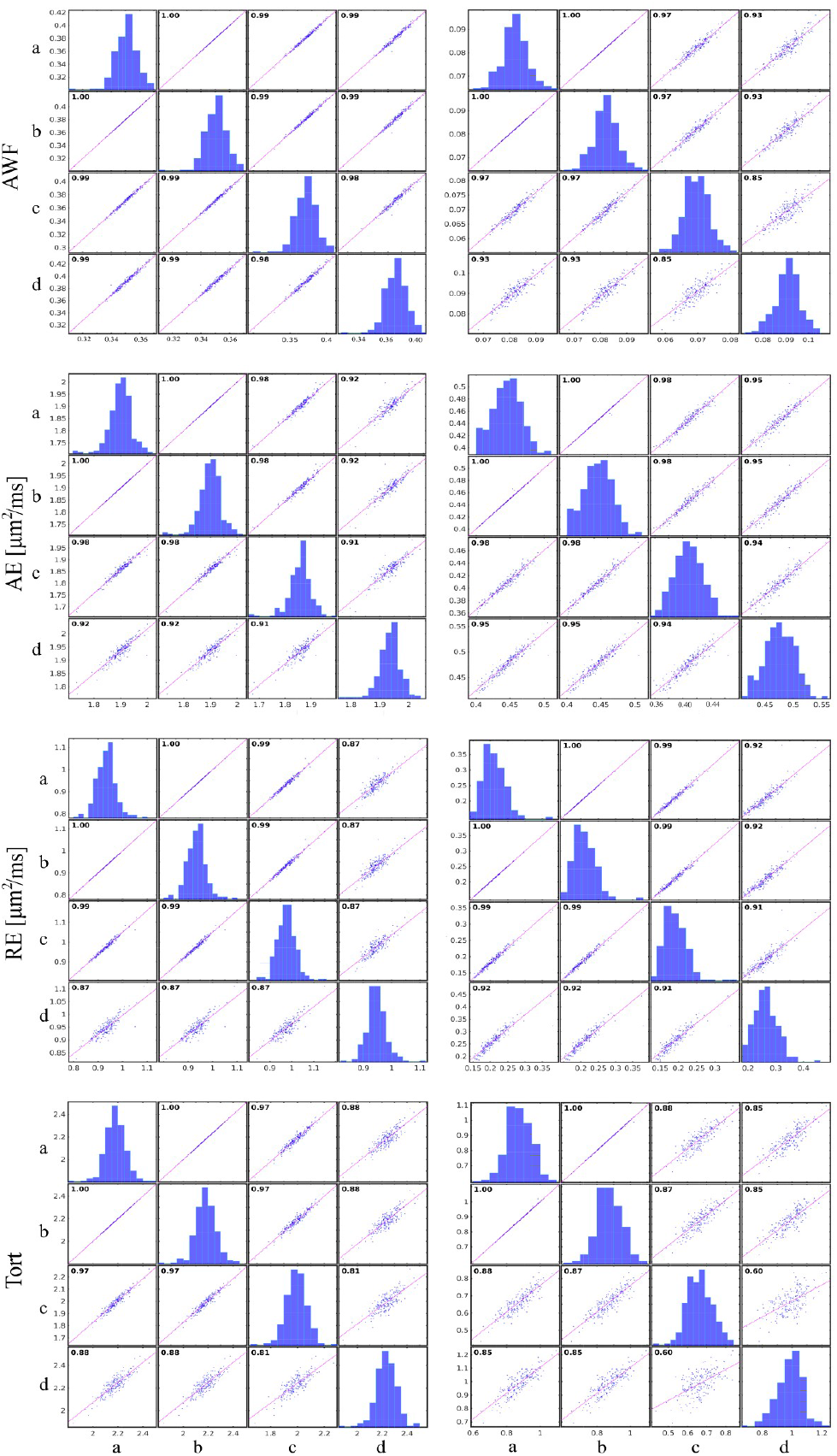
Correlation plots for diffusion metrics based on DKI fitting obtained for four different pipelines: a) up to Step 5; b) up to Step 4; c) up to Step 7; d) original UKB pipeline. Diffusion metrics were averaged over estimated subject skeletons in the case of each pipeline in accordance with the TBSS preparation pipeline.

Figure 6 shows the results from the voxel-wise comparison between the original UKB pipeline and S7. Both DKI/WMTI (Fig. 5a) and cDTI (Fig. 5b) revealed significant differences (p < 0.05) between pipelines, where the S7 pipeline metrics revealed both higher and lower values compared to UKB pipeline. The results of the analysis based on S5 vs S4 and S5 vs S7 are presented in Figure 7.

**Figure 5.**
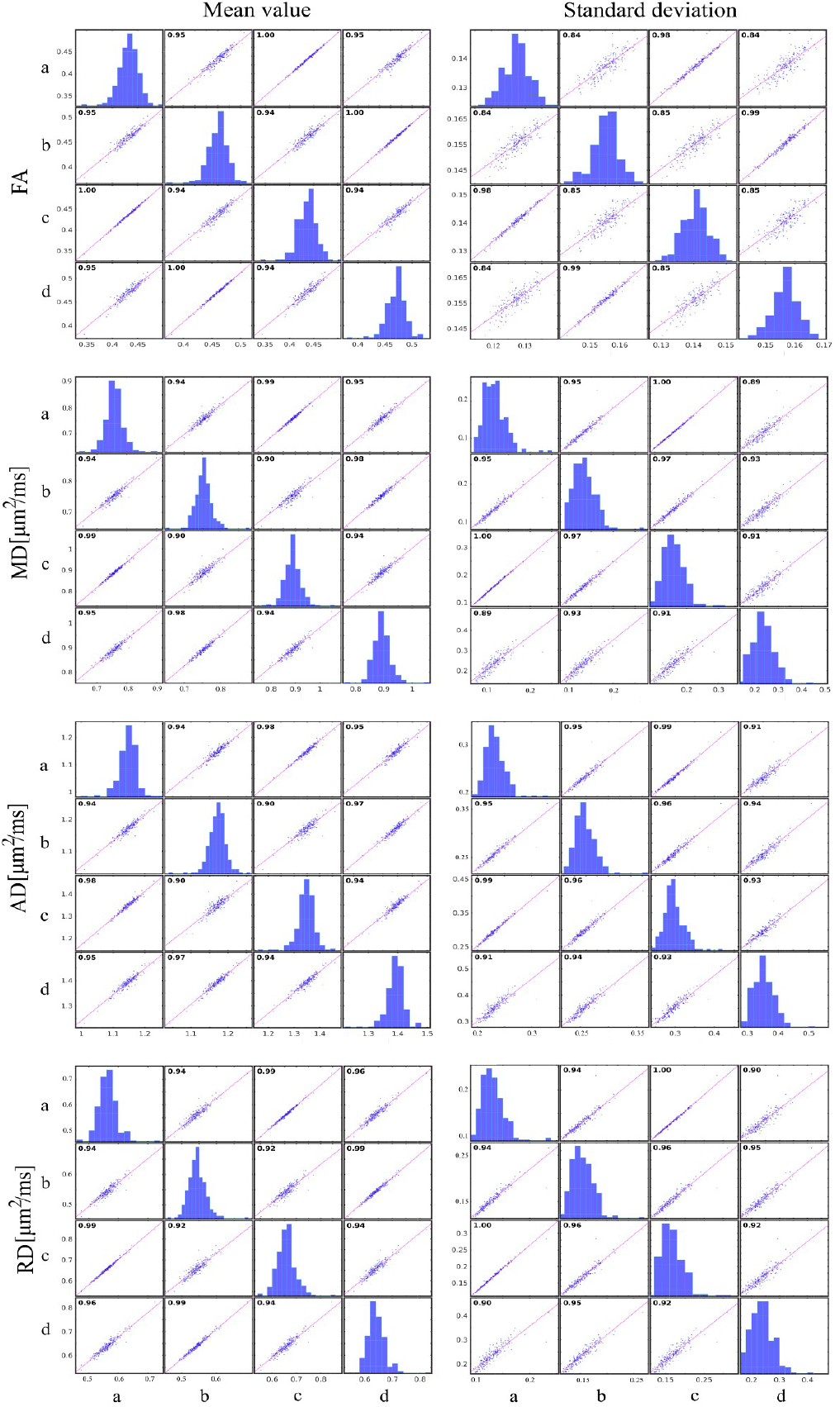
Correlation plots for diffusion metrics based on conventional DTI fitting obtained for four different pipelines: a) up to Step 5; b) up to Step 4; c) up to Step 7; d) original UKB pipeline. Diffusion metrics are averaged over estimated subject skeletons in the case of each pipeline in accordance with TBSS preparation pipeline.

**Figure 6.**
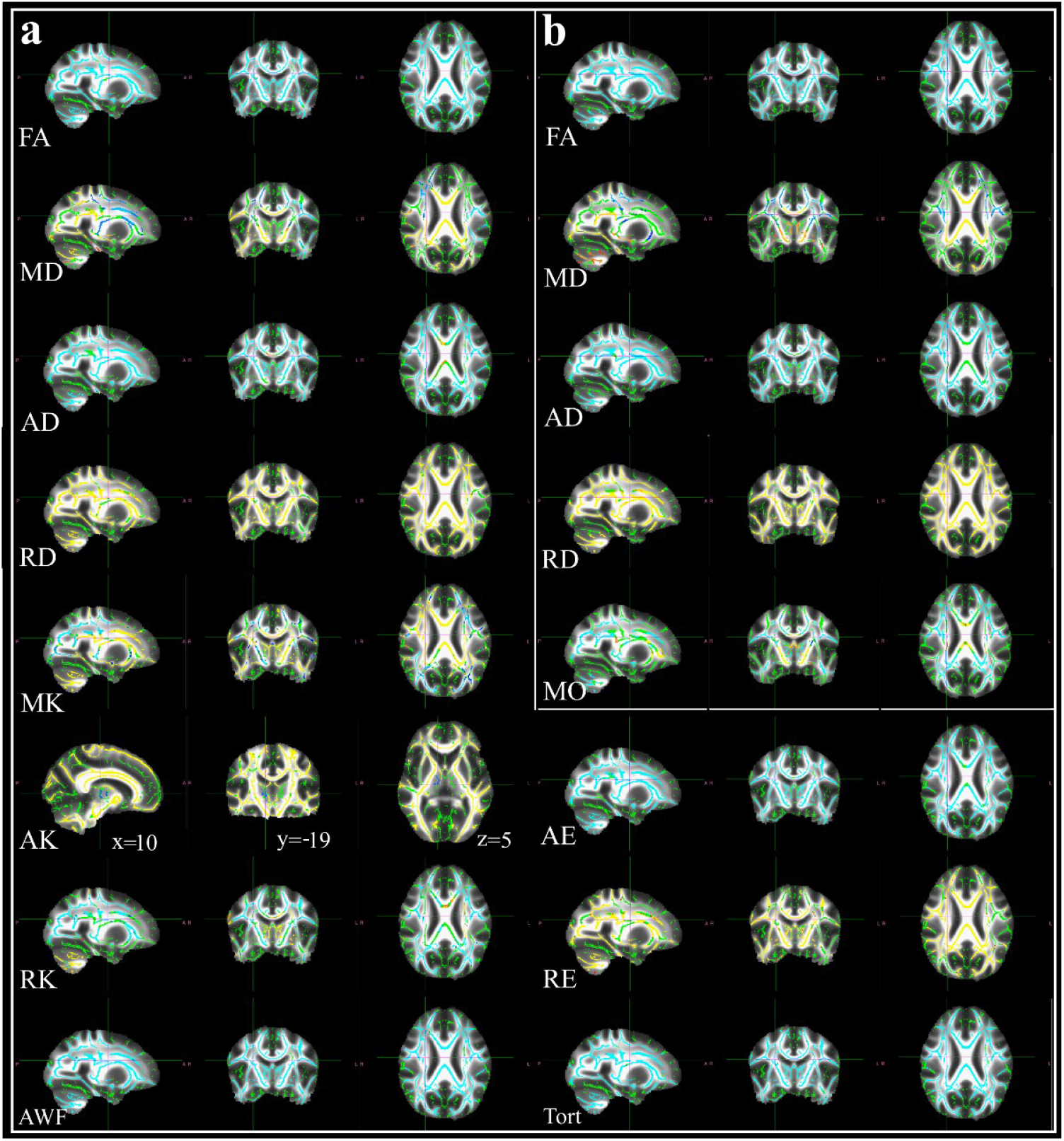
Results of TBSS analysis between the pipelines: original UKB and the proposed here (S7). a) TBSS analysis for diffusion metrics based on DKI and WMTI approaches (2 ?-shells); b) TBSS analysis for diffusion metrics based on DTI fitting (1 *b*-shell). All images are in standard MNI space and correspond to the coordinates: x = 26; y = −8; z = 24. Any coordinate changes are marked. The red-yellow colour means that metrics from S6 pipeline are significantly higher than from UKB (p < 0.05); the blue-light-blue colour means an opposite situation.

**Figure 7.**
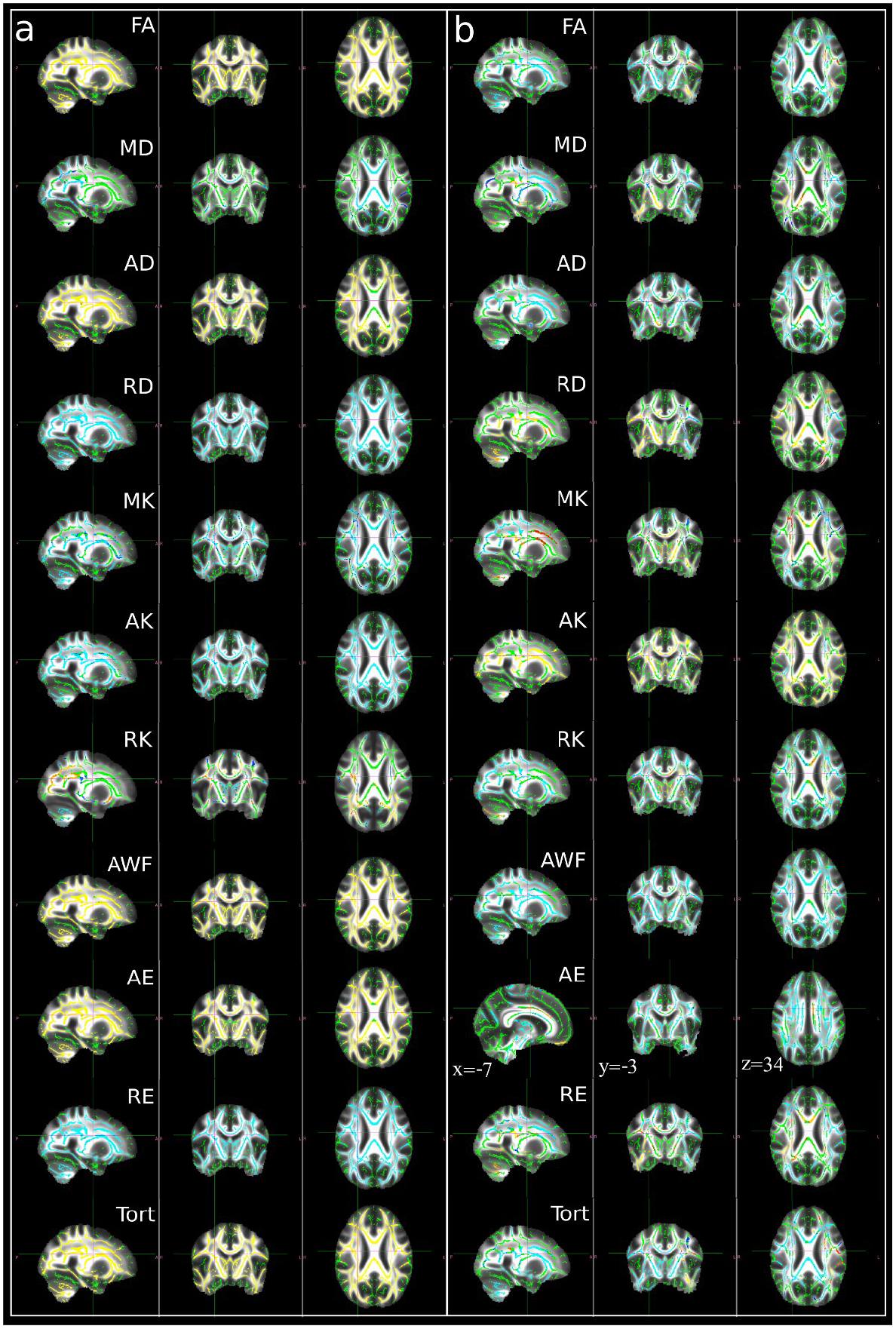
Results of TBSS analysis between a) S5 and S7; b) S5 and original UKB pipeline. All images are in standard MNI space and correspond to the coordinates: x = 26; y = −8; z = 24. Any coordinate changes are marked. The red-yellow colour means that the first pipeline is significantly higher than the second one (p < 0.05); the blue-light-blue colour means an opposite situation.

### tSNR

For *b* = 1000 s/mm^2^ mean estimated tSNR (std) = 3.83 (0.25), 3.83 (0.25), 3.94 (0.29) and 1.81 (0.10) for S4, S5, S7, and UKB, respectively. For *b* = 2000 s/mm^2^ mean estimated tSNR (std) = 1.92 (0.12), 1.92 (0.12), 1.93 (0.14), and 1.73 (0.10). These results indicate 2.2 times (for *b* = 1000 s/mm^2^) and 1.1 time (for *b* = 2000 s/mm^2^) higher tSNR in the S7 pipeline compared to the UKB pipeline.

### Age-related differences across pipelines

Figure 8 shows the estimated age-curves for the various diffusion metrics and Table 2 shows the summary stats from the regression models. The cDTI metrics exhibited expected age-related differences with lower FA and higher MD, AD, and RD with higher age. MK, AK, and RK showed age-related reductions, i.e. reduced non-Gaussianity of the water diffusion with increased age. Metrics based on WMTI (AWF, AE, and RE) demonstrated reduction of the axonal water fraction and extension of the extra-axonal water diffusivity with increased age. Though the main tendency in all age-curves was similar for both pipelines, we found that the Pearson correlations and slopes were different for FA, MK, RK, and AWF, and significantly for AE indicating stronger associations with age in S7 pipeline.

**Figure 8.**
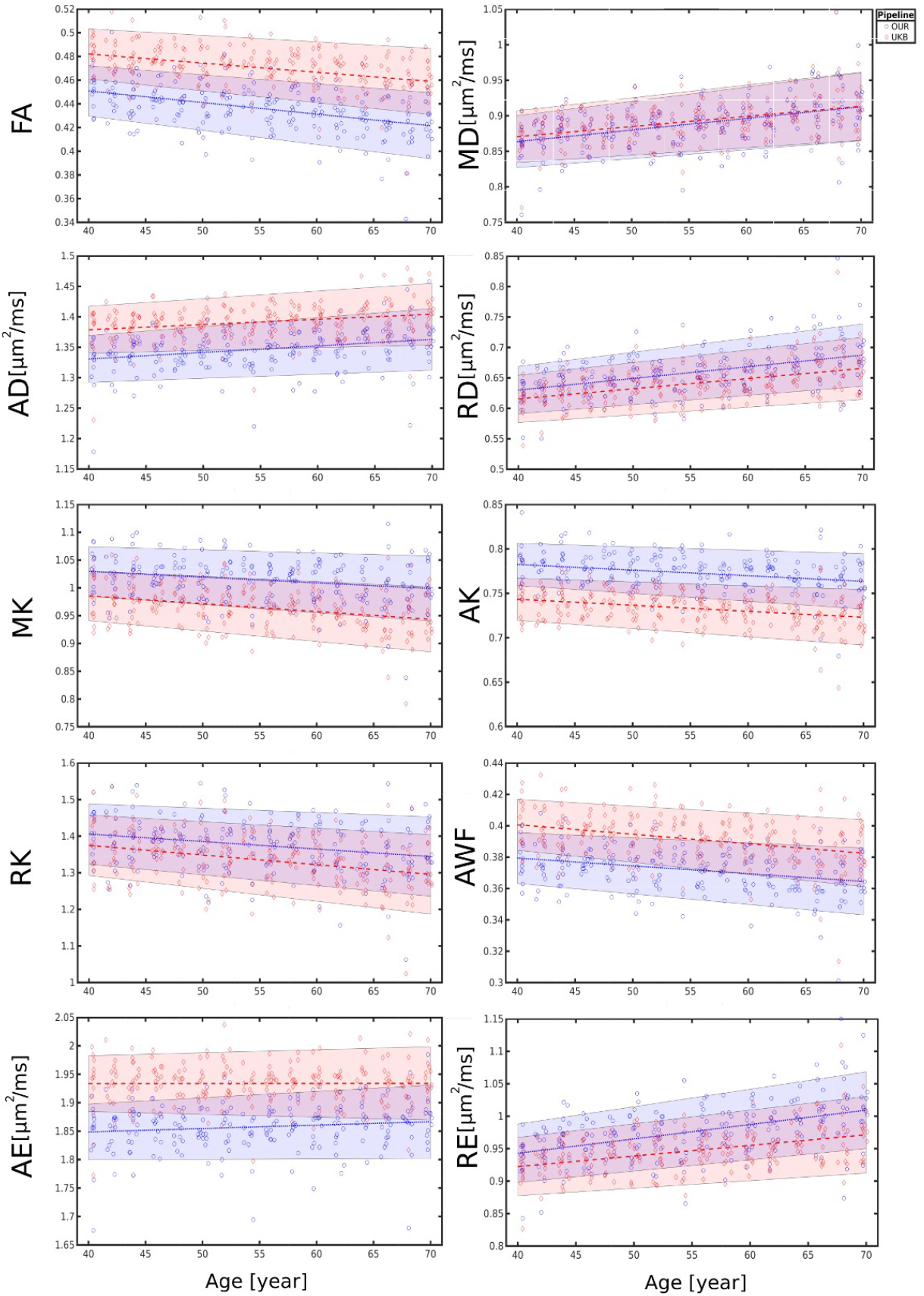
Linear age correlations of diffusion metrics obtained from two pipelines (S7 and original UKB). Colour rectangles present an 95% interval of confidence. The red colour corresponds to UKB values, the blue colour corresponds to the S7 pipeline values.

**Table 2.** Estimated regression slopes/intercepts and Pearson correlations coefficient r/r^2^ for diffusion metrics in age-curves in Fig. 8. The red colour emphasises the significant differences (p < 0.003) in the estimated parameters obtained after applied Fisher’s transformation.

| Pipelines | FA | MD | AD | RD | MK | AK | RK | AWF | AE | RE |
| --- | --- | --- | --- | --- | --- | --- | --- | --- | --- | --- |
| metrics |  |  |  |  |  |  |  |  |  |  |
| **S7** |  |  |  |  |  |  |  |  |  |  |
| *slope* | -1e-3/ | 1.6e-3/ | 1.1e-3/ | 1.9e-3/ | -1e-3/ | -0.6e-3/ | -2.1e-3/ | -0.5e-3/ | 0.6e-3/ | 2.2e-3/ |
| *intercept* | 0.49 | 0.798 | 1.29 | 0.553 | 1.07 | 0.808 | 1.49 | 0.399 | 1.83 | 0.854 |
| **UKB** |  |  |  |  |  |  |  |  |  |  |
| *slope* | -0.8e-3/ | 1.4e-3/ | 0.9e-3/ | 1.7e-3/ | -1.4e-3/ | -0.7e-3/ | -2.7e-3/ | -0.6e-3/ | 0 | 1.6e-3/ |
| *intercept* | 0.513 | 0.813 | 1.35 | 0.549 | 1.04 | 0.771 | 1.48 | 0.425 | 1.93 | 0.857 |
| **S7** |  |  |  |  |  |  |  |  |  |  |
| *r* | -0.44/ | 0.43/ | 0.29/ | 0.47/ | -0.24/ | -0.28/ | -0.26/ | -0.31/ | 0.13/ | 0.47/ |
| *r<sup>2</sup>* | 0.19 | 0.19 | 0.08 | 0.22 | 0.06 | 0.08 | 0.07 | 0.09 | 0.02 | 0.22 |
| **UKB** |  |  |  |  |  |  |  |  |  |  |
| *r* | -0.39/ | 0.44/ | 0.26/ | 0.47/ | -0.33/ | -0.32/ | -0.33/ | -0.35/ | 0.39/ | 0.45/ |
| *r<sup>2</sup>* | 0.15 | 0.19 | 0.07 | 0.22 | 0.11 | 0.10 | 0.11 | 0.12 | 0.15 | 0.20 |

Figure 9 shows the results from general linear model (GLM) testing linear age associations with the different diffusion metrics across the skeleton. The column “Diff” for the same diffusion metrics in Figure 9 visualizes the voxel-wise difference in the correlations. Voxels in red and blue showed significant age association only in S7 and UKB pipelines, respectively. The voxels where both pipelines detected significant age correlations are not shown. The relationship between the number of voxels marked as pipeline specific (red or blue) to the total number of voxels (yellow-red for each pipeline S7 and UKB) is summarised in Table 3. For example, for DTI metrics such as MD and AD the S7 pipeline identified strong age correlations in splenium as well as in the occipital white matter. Similar behaviour is observed for WMTI metrics with strong associations detected by S7 pipeline. In turn, DKI metrics exhibited more regions with strong correlations detected by UKB pipeline.

**Figure 9.**
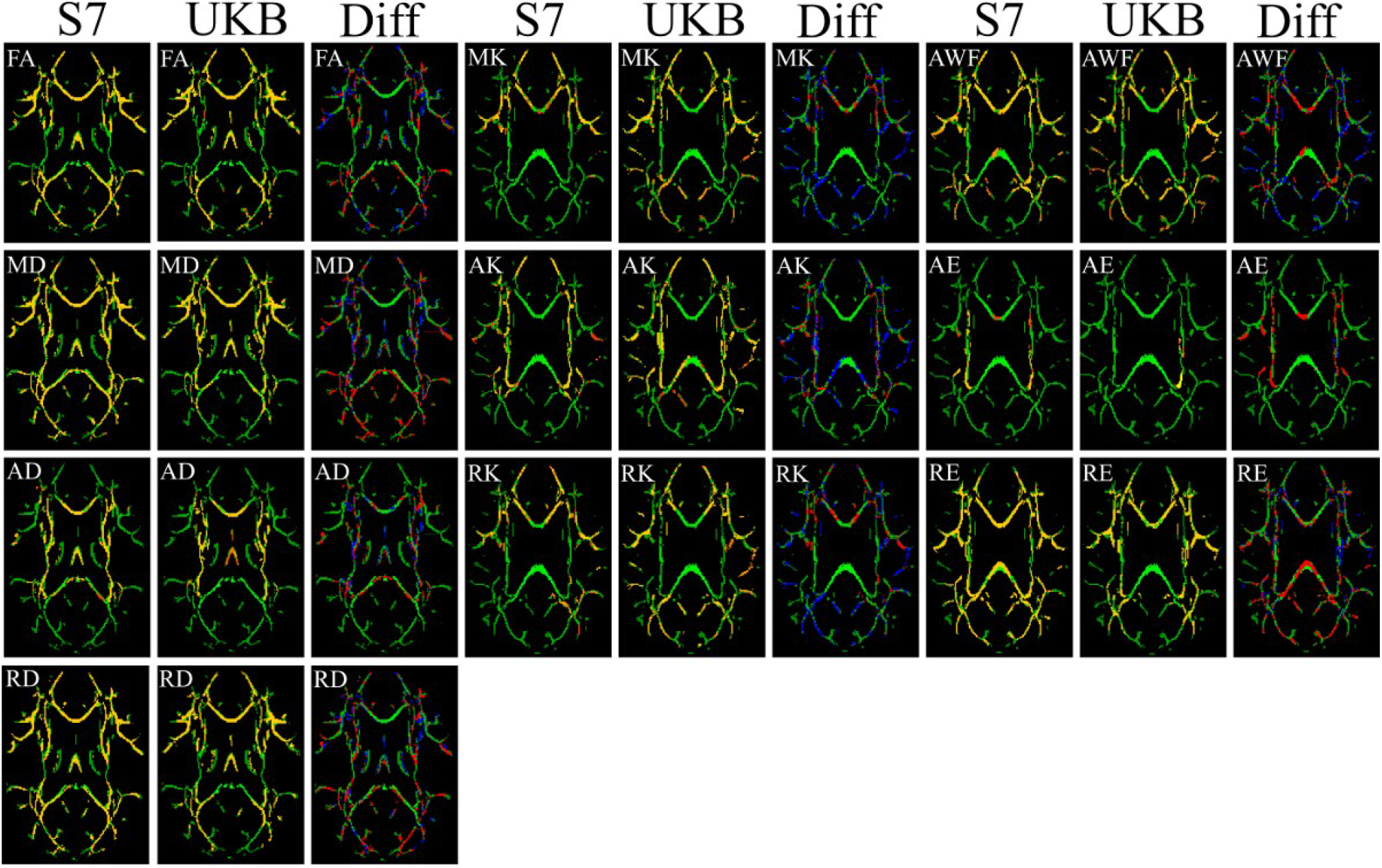
Results of General Linear Model (GLM) tests of diffusion metricses vs age across the skeleton. The linear fitting model is based on two pipelines S7 and UKB. “Diff” columns visualise the spatial difference between the GLM results: the red colour marked the regions with significant difference (p < 0.001) detected along S7 pipeline but not in UKB based tests; the blue colour marked the voxels with significant age correlation detected along UKB pipeline but not in S7. The mean skeleton is visualised by the green colour.

**Table 3.** The number of voxels depending on the pipeline with significant age-correlation specific for the given pipeline vs a total number of voxels with significant age correlation (p < 0.001). See also Fig. 9.

| pipeline<br>specific/total<br>numbers | FA | MD | AD | RD | MK | AK | RK | AWF | AE | RE |
| --- | --- | --- | --- | --- | --- | --- | --- | --- | --- | --- |
| S7 | 35151/<br>70335 | 45335/<br>79974 | 20385/<br>30240 | 43889/<br>87989 | 10816/<br>26896 | 16210/<br>33217 | 18620/<br>39786 | 21452/<br>52974 | 7525/<br>8436 | 47265/<br>81148 |
| UKB | 23744/<br>58928 | 19593/<br>54232 | 7034/<br>16889 | 23717/<br>67817 | 43297/<br>59377 | 27680/<br>44687 | 39003/<br>60169 | 38180/<br>69702 | 832/<br>1743 | 19421/<br>53304 |

## Discussion

A growing interest in utilising advanced diffusion weighted imaging to study the human brain motivated us to test the effects of various data processing pipelines on different diffusion metrics. Differences in data post-processing steps are likely to influence reliability and results. Thus, a harmonised diffusion pipeline may prove valuable for increasing sensitivity, reliability, and generalisability across studies. We suggest a general framework with the following post-processing steps: 1) noise correction, 2) Gibbs ringing correction, 3) field mapping, 4) susceptibility, eddy current and motion distortion corrections, 5) B1 field correction, 6) spatial smoothing; and 7) final metrics estimation. Our comparison between four diffusion pipelines demonstrated that the general pipeline suggested here yield a substantially higher tSNR compared to the original UKB pipeline, and also influence the estimated age curves for MK, RK, AE, and AWF, with potentially important implications for the interpretations.

Overall, the diffusion metrics derived using the different pipelines demonstrated high correlations and similar standard deviations. In some cases, S7 resulted in lower standard deviations of the diffusion metrics than others, e.g. for WMTI. S7 also exhibited high correlations with S5 for the conventional DTI metrics. Results from the UKB pipeline showed relatively high correlations with S4 and S5 for the DTI metrics, and slightly lower for DKI and WMTI. For cDTI the UBK pipeline showed stronger correlations with S4 than to S5. The correlations between the S7 and UKB pipelines were lower than for other pipeline pairs. Overall, the results support that all blocks of the proposed S7 pipeline might lead to relevant improvements in the estimations of absolute diffusion metrics.

TBSS revealed a significant difference between the S7 and UKB pipelines for all diffusion metrics. Interestingly, the differences between pipelines did not reflect global shifts of the diffusion parameters across the skeleton, but rather spatially variable differences across several metrics, including DTI, DKI, and WMTI. The observed differences between S4 and S5 pipelines suggest significant effects of bias field corrections across a large part of the skeleton, nevertheless, the difference between the S4 and S5 pipelines looks similar to the value shift in contrast to the case of S7 and UKB comparison. The comparison between S5 and UKB pipelines revealed similar results as those seen when comparing the S7 and UKB pipelines (Fig 7b), suggesting that spatial smoothing in the S7 pipeline is a reasonable improvement which did not remove important information from the dataset. The findings of regionally specific increases or decreases in diffusion metrics for different pipelines might partly be explained by an effect of the noise correction step on physiological noise around the large arteries or strong susceptibility artefacts close to air cavities in the brain. Such artefacts might introduce spatially variable distortions, which could lead to spurious findings in clinical studies. This could explain the previously demonstrated higher sensitivity to group differences after noise correction (Kochunov et al., 2018). Moreover, our comparison between pipelines demonstrated that noise and Gibbs ringing corrections influenced tSNR both in the case of conventional diffusion data *(b* = 1000 s/mm^2^) and at higher diffusion weightings *(b* = 2000 s/mm^2^). In contrast, the spatial smoothing step (S7) did not introduce a strong shift in tSNR in our analysis (S4, S5), however, has been shown to influence further diffusion metric estimations by reducing the number of “bad” voxels (Veraart et al., 2013).

In order to assess possible practical consequences of the different noise correction steps we compared the estimated age slopes in DKI and WMTI metrics between pipelines. Age-related differences are abundant in the relevant age span (Grinberg et al., 2017; Westlye et al., 2010). Apart from MD and RD, all included metrics showed a large main effect of pipeline and, although similar signs, different estimated age-slopes between the S7 and UKB pipelines. In particular, the S7 pipeline revealed stronger age-correlations compared to UKB for WMTI metrics. Further, the voxel-wise comparisons revealed a higher number of voxels showing significant age associations in the S7 compared to the UKB pipeline for DTI and WMTI metrics. On the contrary, the UKB pipeline demonstrated a higher number of significant voxels in the case of DKI estimations. Although subtle, pipeline related global and spatially varying differences in diffusion metrics will have consequences for subsequent analyses, for example, for machine-learning based age prediction or diagnostic classification or prediction of clinical traits (Alnaes et al., 2018; Doan et al., 2017; Kuhn at al., 2018; Richards et al., 2018).

In conclusion, our analysis demonstrated that diffusion metric estimations are sensitive to specific pipeline and might benefit from the proposed sequential advanced post-processing steps. Our proposed pipeline is also an example of a general approach for harmonisation of postprocessing steps across diffusion MRI studies for increased sensitivity and generalisability, which may be particular important when applying complex diffusion models to multishell diffusion data.

## Acknowledgement

This work was funded by the Research Council of Norway (249795) and the South-Eastern Norway Regional Health Authority (2014097, 2015073, 2016083).

## Conflict of interests

Authors declared no conflict of interests.

## Supplementary Materials

An example of diffusion bash scripts used in the work are accessible as a text file.

## References

Aja-Fernandez S., Vegas-Sanchez-Ferrero G., Tristan-Vega A., 2014. Noise estimation in parallel MRI: GRAPPA and SENSE. Magn. Reson. Imaging 32, 281–290.

Alfaro-Almagro F., Jenkinson M., Bangerter N.K., Andersson J.L.R., et al., 2018. Image processing and Quality Control for the first 10000 brain imaging datasets from UK Biobank. Neuroimage 166, 400–424.

Alnaes D., Kaufmann T., Doan N.T., Cordova-Palomera A., Wang Y., Bettella F., Moberget T., Andreassen O.A., Westlye L.T., 2018. Association of Heritable Cognitive Ability and Psychopathology With White Matter Properties in Children and Adolescents. JAMA Psychiatry 75, 287–295.

Andersson J.L.R., Skare S., Ashburner J., 2003. How to correct susceptibility distortions in spinecho echo-planar images: application to diffusion tensor imaging. Neuroimage, 20, 870–888.

Andersson J.L.R., Jenkinson M., Smith S., 2007. Non-linear optimisation In: FMRIB technical report TR07JA1 from www.fmrib.ox.ac.uk/analysis/techrep

Andersson J.L.R., Sotiropoulos S.N., 2016a. An integrated approach to correction for off-resonance effect and subject movement in diffusion MR imaging. Neuroimage 125, 1063–1078.

Andersson J.L.R., Graham M.S., Zsoldos E., Sotiropoulos S.N., 2016b. Incorporating outlier detection and replacement into a non-parametric framework for movement and distortion correction of diffusion MR imaging. Neuroimage 141, 556–572.

Andersson J.L.R., Graham M.S., Drobnjak I., Zhang H., Filippini, Bastiani M., 2017. Towards a comprehensive framework for movement and distortion correction of diffusion MR imaging: Within volume movement. Neuroimage 152, 450–466.

Andre E.D., Grinberg F., Farrher E., Maximov I.I., Shah N.J., Meyer C., Jaspar M., Muto V., Phillips C., Balteau E., 2014. Influence of noise correction on intra- and inter-subject variability of quantitative metrics in diffusion kurtosis imaging. Plos One 9, e94531.

Banerjee A., Maji P., 2015. Rough sets and stomped normal distribution for simultaneous segmentation and bias field correction in brain MR images. IEEE Image Processing 24, 5764–5776.

Basser P.J., Mattiello J., Le Bihan D., 1994. MR diffusion tensor spectroscopy and imaging. Bioph. J. 66, 259–267.

Bastiani M., Andersson J.L.R., Cordero-Grande L., Murgasova M., Hutter J., Price A.N., Makropoulos A., Fitzgibbon S.P., Hughes E., Rueckert D., Victor S., Rutherford M., Edwards A.D., Smith S.M., Tournier J.D., Hajnal J.V., Jbabdi S., Sotiropoulos S.N., 2019. Automated processing pipeline for neonatal diffusion MRI in the developing Human Connectome Project. Neuroimage 185, 750–763.

Ciu Z., Zhong S., Xu P., He Y., Gong G., 2013. PANDA: a pipeline toolbox for analyzing brain diffusion images. Frontiers in Human Neuroscience 7, 42.

Doan N.T., Persson A.E.K., Alnaes D., Kaufmann T., Rokicki J., Cordova-Palomera A., Moberget T., Braekhus A., Barca M.L., Engedal K., Andreassen O.A., Selbaek G., Westlye L.T., 2017. Dissociable diffusion MRI patterns of white matter microstructure and connectivity in Alzheimer's disease spectrum. Scientific Reports 7, 45131.

Esteban O., Birman D., Schaer M., Koyejo O.O., Poldrack R.A., Gorgolewski K.J., 2017. MRIQC: Advancing the automatic prediction of image quality in MRI from unseen sites. PLOS ONE 12, e0184661.

Farzinfar M., Oguz I., Smith R.G., Verde A.R., Dietrich C., Gupta A., Escolar M.L., Piven J., Pujol S., Vachet C., Gouttard S., Gerig G., Dager S., McKinstry R.C., Paterson S., Evans A.C., The IBIS network, Styner M.A., 2013. Diffusion imaging quality control via entropy of principal direction distribution. Neuroimage 82, 1–12.

Fieremans E., Jensen J.H., Helpern J.A., 2011. White matter characterization with diffusion kurtosis imaging. Neuroimage 58, 177–188.

Gallichan D., Scholz J., Bartsch A., Behrens T.E., Robson M.W., Miller K.L., 2010. Addressing a systematic artifact in diffusion-weighted MRI. Human Brain Mapping 31, 193–202.

Ganzetti M., Wenderoth N., Mantini D., 2016. Quantitative evaluation of intensity inhomogeneity correction methods for structural MR brain images. Neuroinformatics 14, 5–21.

Glasser M.F., Sotiropoulos S.N., Wilson J.A., Coalson T.S., Fischl B., Andersson J.L., Xu J., Jbabdi S., Webster M., Polimeni J.R., Van Essen D.C., Jenkinson M., for the WU-Minn HCP Consortium, 2013. The minimal preprocessing pipelines for the Human Connectome Project. Neuroimage 80, 105–124.

Grinberg F., Maximov I.I., Farrher E., Neuner I., Amort L., Thoennessen H., Oberwelland E., Konrad K., Shah N.J., 2017. Diffusion kurtosis metrics as biomarkers of microstructural development: A comparative study of a group of children and a group of adults. Neuroimage 144 (Pt. A), 12–22.

Hasan K.M., 2007. A framework for quality control and parameter optimization in diffusion tensor imaging: theoretical analysis and validation. Magn. Reson. Imaging 25, 1196–1202.

Jensen J.H., Helpern J.A., Ramani A., Lu H., Kaczynski K., 2005. Diffusion kurtosis imaging: the quantification of non-gaussian water diffusion by means of magnetic resonance imaging. Magn. Reson. Med. 53, 1432–1440.

Johansen-Berg H., Behrens T.E.J., 2014. Diffusion MRI: From quantitative measurement to in-vivo neuroanatomy. Elsevier Academic Press.

Kellner E., Dhital B., Kiselev V.G., Reisert M., 2016. Gibbs-ringing artifact removal based on local subvoxel-shifts. Magn. Reson. Med. 76, 1574–1581.

Kochunov P., Dickie E.W., Viviano J.D., Turner J., Kingley P.B., Jahanshad N., Thompson P.M., Ryan M.C., Fieremans E., Novikov D.S., Veraart J., Hong E.L., Malhotra A.K., Buchanan R.W., Chavez S., Voineskas V.N., 2018. Integration of routine QA data into meta-analysis may improve quality and sensitivity of multisite diffusion tensor imaging studies. Human Brain Mapping 39, 1015–1023.

Kuhn T., Kaufmann T., Doan N.T., Westlye L.T., Jones J., Nunez R.A., Bookheimer S.Y., Singer E.J., Hinkin C.H., Thames A.D., 2018. An augmented aging process in brain white matter in HIV. Human Brain Mapping 39, 2532–2540.

Maximov I.I., Grinberg F., Shah N.J., 2011. Robust tensor estimation in diffusion tensor imaging. J. Magn. Reson. 213, 136–144.

Maximov I.I., Farrher E., Grinberg F., Shah N.J., 2012. Spatially variable Rician noise in magnetic resonance imaging. Medical Image Analysis 16, 536–548.

Maximov I.I., Thoennessen H., Konrad K., Amort L., Neuner I., Shah N.J., 2015. Statistical instability of TBSS analysis based on DTI fitting algorithm. J. Neuroimaging 25, 883–891.

Maximov I.I., Tonoyan A.S., Pronin I.N., 2017. Differentiation of glioma malignancy grade using diffusion MRI. Physica Medica 40, 24–32.

McRobbie D.W., Moore E.A., Graves M.J., Prince M.R., 2009. MRI from Picture to Proton. Cambridge University Press.

Miller K.L., Alfaro-Almagro F., Bangerter N.K., Thomas D.L., Yacoub E., Xu J., Bartsch A.J., Jbabdi S., Sotiropoulos S.N., Andersson J.L., Griffanti L., Douaud G., Okell T.W., Weale P., Dragonu I., Garratt S., Hudson S., Collins R., Jenkinson M., Matthews P.M., Smith S.M., 2016. Multimodal population brain imaging in the UK Biobank prospective epidemiological study. Nature Neuroscience 19, 1523–1536.

Novikov D.S., Kiselev V.G., Jespersen S.N., 2018. On modeling. Magnetic Resonance in Medicine 79, 3172–3193.

Oguz I., Farzinfar M., Matsui J., Budin F., Liu Z., Gerig G., Johnson H.J., Styner M., 2014. DTIPrep: quality control of diffusion-weighted images. Frontiers in Neuroinformatics 8, 4.

Perona P., Malik J., 1987. Scale-space and edge detection using anisotropic diffusion. Proc. IEEE Comp. Soc. Workshop Comp. Vision 16–22.

Perrone D., Aelterman J., Puzurica A., Jeurissen B., Philips W., Leemans A., 2015. The effect of Gibbs ringing artifacts on measures derived from diffusion MRI. Neuroimage 120, 441–455.

Richard G., Kolskaar K., Sanders A.M., Kaufmann T., Petersen A., Doan N.T., Sanchez J.M., Alnaes D., Ulrichsen K.M., Dorum E.S., Andreassen O., Nordvik J.E., Westlye L.T., 2018. Assessing distinct patterns of cognitive aging using tissue-specific brain age prediction based on diffusion tensor imaging and brain morphometry. BioRxiv doi: https://doi.org/10.1101/313015.

Roalf D.R., Quarmley M., Elliott M.A., Satterthwaite T.D., Vandekar S.N., Ruparel K., Gennatas E.D., Calkins M.E., Moore T.M., Hopson R., Prabhakaran K., Jackson C.T., Verma R., Hakonarson H., Gur R.C., Gur R.E., 2016. The impact of quality assurance assessment on diffusion tensor imaging outcomes in a large-scale population-based cohort. Neuroimage 125, 903–319.

Sairanen V., Leemans A., Tax C.M.W., 2018. Fast and accurate slisewise outlier detection (SOLID) with informed model estimation for diffusion MRI data. Neuroimage 181, 331–346.

Smith S.M., Jenkinson M., Woolrich M.W., Beckmann C.F., Behrens T.E.J., Johansen-Berg H., Bannister P.R., De Luca M., Drobnjak I., Flitney D.E., Niazy R., Saunders J., Vickers J., Zhang Y., De Stefano N., Brady J.M., Matthews P.M., 2004. Advances in functional and structural MR image analysis and implementation as FSL. Neuroimage, 23, 208–219.

Smith S.M. Jenkinson M., Johansen-Berg H., Rueckert D., Nichols T.E., Mackay C.E., Watkins K.E., Ciccarelli O., Cader M.Z., Matthews P.M., Behrens T.E., 2006. Tract-based spatial statistics: voxelwise analysis of multi-subject diffusion data. Neuroimage 31, 1487–1505.

Smith S.M., Johansen-Berg H., Jenkinson M., Rueckert D., Nichols T.E., Miller K.L., Robson M.D., Jones D.K., Klein J.C., Bartsch A.J., Behrens T.E., 2007. Acquisition and voxelwise analysis of multi-subject diffusion data with tract-based spatial statistics. Nat. Protoc. 2, 499–503.

Smith S.M. Nichols T.E., 2009. Threshold-free cluster enhancement: addressing problems of smoothing, threshold dependence and localisation in cluster inference. Neuroimage 44, 83–98.

Sotiropoulos S.N., Jbabdi S., Xu J., Andersson J.L., Moeller S., Auerbach E.J., Glasser M.F., Hernandez M., Sapiro G., Jenkinson M., Feinberg D.A., Yacoub E., Lenglet C., Van Essen D.C., Ugurbil K., Behrens T.E.J., for the WU-Minn HCP Consortium, 2013. Advances in diffusion MRI acquisition and processing in the Human Connectome Project. Neuroimage 80, 125–143.

Tamnes C.K., Roalf D.R., Goddings A.L., Lebel C., 2017. Diffusion MRI of white matter microstructure development in childhood and adolescence: Methods, challenges and progress. Dev. Cong. Neurosci. doi: 10.1016/j.dcn.2017.12.002.

Taylor P.A., Alhamud A., van der Kouwe A., Saleh M.G., Laughton B., Meintjes E., 2016. Assessing the performance of different DTI motion correction strategies in the presence of EPI distortion correction. Human Brain Mapping 37, 4405–4424.

Tønnesen S., Kaufmann T., Doan N.T., Alnæs D., Córdova-Palomera A., Meer D.V., Rokicki J., Moberget T., Gurholt T.P., Haukvik U.K., Ueland T., Lagerberg T.V., Agartz I., Andreassen O.A., Westlye L.T., 2018. White matter aberrations and age-related trajectories in patients with schizophrenia and bipolar disorder revealed by diffusion tensor imaging. Sci. Rep. 8, 14129.

Tustison N.J., Avants B.B., Cook P.A., Zheng Y., Egan A., Yushkevich P.A., Gee J.C., 2010. N4ITK: improved N3 bias correction. IEEE Trans. Med. Imaging 29, 1310–1320.

Van Hecke W., Leemans A., De Backer S., Jeurissen B., Parizel P.M., Sijbers J., 2010. Comparing isotropic and anisotropic smoothing for voxel-based DTI analysis: a simulation study. Human Brain Mapping 31, 98–114.

Vellmer S., Tonoyan A.S., Suter D., Pronin I.N., Maximov I.I., 2018. Validation of DWI preprocessing procedures for reliable differentiation between human brain gliomas. Z. Med. Phys. 28, 14–24.

Veraart J., Sijbers J., Sunaert S., Leemans A., Jeurissen B., 2013. Weighted linear least squares estimation of diffusion MRI parameters: strenghts, limitations, and pitfalls. Neuroimage 81, 335–346.

Veraart J., Fieremans E., Novikov D.S., 2016a. Diffusion MRI noise mapping using random matrix theory. Magn. Reson. Med. 76, 1582–1593.

Veraart J., Novikov D.S., Christiaens D., Ades-Aron B., Sijbers J., Fieremans E., 2016b. Denoising of diffusion MRI using random matrix theory. Neuroimage 142, 394–406.

Veraart J., Fieremans E., Jelescu I.O., Knoll F., Novikov D.S., 2016c. Gibbs ringing in diffusion MRI. Magn. Reson. Med. 76, 301–314.

Walker L., Chang L.C., Koay C.G., Sharma N., Cohen L., Verma R., Pierpaoli C., 2011. Effect of physiological noise in population analysis of diffusion tensor MRI data. Neuroimage 54, 1168–1177.

Westlye L.T., Walhovd K.B., Dale A.M., Bjørnerud A., Due-Tønnessen P., Engvig A., Grydeland H., Tamnes C.K., Ostby Y., Fjell A.M., 2010. Life-span changes of the human brain white matter: diffusion tensor imaging (DTI) and volumetry. Cereb. Cortex 20, 2055–2068.

Westlye L.T., Reinvang I., Rootwelt H. Espeseth T., 2012. Effects of APOE on brain white matter microstructure in healthy adults. Neurology 79, 1961–1969.

Winkler A.M., Ridgway G.R., Webster M.A., Smith S.M., Nichols T.E., 2014. Permutation inference for the general linear model. Neuroimage 92, 381–297.

Wu M., Chang L.C., Walker L., Lemaitre H., Barnett A.S., Marenco S., Pierpaoli C., 2008. Comparison of EPI distortion correction methods in diffusion tensor MRI using a novel framework. Med. Image Comput. Comput. Assist. Interv. 11, 321–329.

